# The specialization of *Esteya vermicola* hyphae in infection to *Bursaphelenchus xylophilus* and its colonization of pine tree

**DOI:** 10.1101/2020.08.21.260729

**Authors:** Hai-Hua Wang, Can Yin, Jie Gao, Ran Tao, Piao-Piao Dai, Chun-Yan Wang, Chang-Keun Sung

## Abstract

Pine wilt disease (PWD) caused by the nematode *Bursaphelenchus xylophilus* is a serious problem on pines, and there is currently no effective control strategy for this disease. Although the endoparasitic fungus *Esteya vermicola* showed great effectiveness in controlling pine wilt disease, the colonization patterns of the host pine tree xylem by this fungus are unknown. To investigate the colonization patterns of pine xylem by this fungus, the species *Pinus koraiensis* grown in a greenhouse was used as an experimental host tree. The fungal colonization of healthy and wilting pine trees by *E. vermicola* was quantified using PCR with a TaqMan probe, and a green fluorescence protein (GFP) transformant was used for visualization. The results reported a specific infection approach used by *E. vermicola* to infect *B. xylophilus* and specialized fungal parasitic cells in PWN infection. In addition, the inoculated blastospores of *E. vermicola* germinated and grew inside of healthy pine xylem, although the growth rate was slow. Moreover, *E. vermicola* extended into the pine xylem following spray inoculation of wounded pine seedling stems, and a significant increase in fungal quantity was observed in response to *B. xylophilus* invasion. An accelerated extension of *E. vermicola* colonization was shown in PWN-infected wilting pine trees, due to the immigration of fungal-infected PWNs. Our results provide helpful knowledge about the extension rate of this fungus in healthy and wilting PWN-susceptible pine trees in the biological control of PWD and will contribute to the development of a management method for PWD control in the field.

**Author summary:** Pine wilt disease, caused by *Bursaphelenchus xylophilus*, has infected most pine forests in Asian and European forests and led to enormous losses of forest ecosystem and economy. *Esteya vermicola* is a bio-control fungus against pinewood nematode, showed excellent control efficient to pine wilt disease in both of greenhouse experiments and field tests. Although this bio-control agent was well known for the management of pine wilt disease, the infection mechanism of fungal infection and colonization of host pine tree are less understand. Here, we use GFP-tagged mutant to investigate the fungal infection to pinewood nematode; additionally, the temporal and spatial dynamics of *E. vermicola* colonize to pine tree were determined by the TaqMan real-time PCR quantification, as well as the response to pinewood nematode invasion. We found a specific infection approach used by *E. vermicola* to infect *B. xylophilus* and specialized fungal parasitic cells in PWN infection. In addition, the fungal germination and extension inside of pine tree xylem after inoculation were revealed. In addition, the quantity of *E. vermicola* increased as response to pinewood nematode invasion was reported. Our study provides two novel technologies for the visualization and detection of *E. vermicola* for the future investigations of fungal colonization and its parasitism against pinewood nematode, and the mechanisms of the bio-control process.

## Introduction

The pinewood nematode (PWN), *Bursaphelenchus xylophilus*, infects pine trees as the causal agent of pine wilt disease (PWD) [1] and is transmitted through the wounds created by the sawyer beetle (*Monochamus* spp.) feeding [2]. The PWN feeds on the epithelial cells and resin ducts of its host, which leads to dysfunction of vascular organs. Subsequently, the PWN is distributed throughout the sapwood of the branches, trunk, and roots of susceptible host plants [3]. Movement of PWNs and vascular dysfunction resulting in wilt and death of host pine trees occur rapidly. The PWN is supposed to be one of the native species in North America. Then, the international trade of timber introduced *B. xylophilus* into Japan, from where it spread to China, Korea, and Europe rapidly. PWD is a striking example of how ecosystems are threatened by the establishment of an exotic organism.

Control strategies for PWD include plant quarantine, integrated control of the vector, silvicultural measures, and replanting with resistant species. A large number of new forest infections occur every year, despite the dedicated and concerted actions of government agencies. *Esteya vermicola* is the first recorded endoparasitic fungus of PWN [4]. The *E. vermicola* strains were isolated from infected PWNs, soil, or dead trees [4-7]. It is well known that *E. vermicola* has high infectivity against PWN, and the species therefore has been widely used as a biological control agent against PWD in the field [8]. Hyphae of *E. vermicola* and parasitized PWNs were observed only from wood sections of dead and wilting seedlings inoculated with a conidial suspension of the fungus [9,10]. Recently, the fungal germination and colonization of *E. vermicola* were live-cell visualized with the help of a green fluorescence protein (GFP)-transformed mutant *CNU120806gfp* [11].

Recent studies documenting colonization by plant-endophytic fungi, such as mycorrhizal fungi, root pathogenic fungi, and soil-inhabiting fungi, were conducted on roots using a range of visualization technologies [12-15]. However, the colonization and development patterns of *E. vermicola* in the host tree remain less known, as well as the fungal responses to PWN invasion in the host plant. The expression of the *gfp* gene greatly helps to improve the knowledge of *E. vermicola* colonization of host plants and *in vivo* infection to PWNs.

Polymerase chain reaction (PCR) amplification has been applied for the molecular detection of various organisms. To detect *E. vermicola* after inoculation or spraying on pine trees, Wei et al. (2014) [16] developed a simple and easy method with FTA-DNA extraction and PCR amplification from environmental samples based on the presence of a specific 176 bp fragment. Subsequently, a novel pinewood sample preparation and DNA extraction method were established [17], in which another three primer pairs based on chitinase, β-tubulin, and large subunit ribosomal RNA genes were designed for PCR amplification. However, the above two techniques cannot provide quantification analyses to reveal the colonization patterns of *E. vermicola*. More recently, a precise and accurate quantitative technique for *E. vermicola* in environmental samples was developed with TaqMan PCR quantification [18].

The fungus was inoculated into pine tree xylem as a nonnative species due to its parasitic behavior against PWNs. Thus, the colonization and development of *E. vermicola* in host pine trees before or during PWN invasion are the most concerning. However, few studies on the temporal and spatial dynamics of fungal colonization of susceptible tree xylem following artificial inoculation has been conducted. Thus, the above-developed approaches were used in the present study to reveal the temporal and spatial patterns of pine xylem colonization by *E. vermicola*, with or without the presence of PWN.

## 2 Results

### 2.1 Construction of calibration curves

In the nested-based TaqMan PCR, the 473-bp and 213-bp DNA products were generated using primers 28S-1F/28S-1R and 28S-2F/28S-2R, respectively. The application of nested TaqMan PCR allowed the detection of target DNA at quantities as low as 10^−5^ ng. The standard curves for real-time TaqMan PCR quantification of both *wt E. vermicola* and *CNU120806gfp* obtained in the present study are shown in Supplementary Fig. S1. The curves showed good fits, with R^2^ values of 0.9838 and 0.9969 for traditional and nested TaqMan PCR of *wt E. vermicola*, respectively (Supplementary Fig. S1a - b). The *C*_*t*_ values of traditional TaqMan PCR ranged from 25.81 ± 0.82 to 41.24 ± 1.04, and those of nested TaqMan PCR ranged from 14.86 ± 0.16 to 32.93 ± 0.07. For *CNU120806gfp*, R^2^ values of 0.9759 and 0.9927 for traditional and nested TaqMan PCR, respectively, were obtained (Supplementary Fig. S1c - d). The detectable concentration of the genomic DNA of pure cultured *E. vermicola* with conventional TaqMan PCR ranged from 10^−2^ ng/μl to 10^2^ ng/μl. However, the lowest detectable concentration of template using nested PCR, which significantly increased the detection sensitivity, was 10^−5^ ng/μl.

### 2.2 *In vitro* colonization patterns of PWN by *E. vermicola*

#### 2.2.1 Temporal infection dynamics of PWN by *E. vermicola*

The fungal infection pattern of PWN by the nematophagous fungus *E. vermicola* was studied by quantification of fungal hyphae. The single-PWNs that were classified by the infection stage were assayed, the infection status of each stage of these tested PWNs is shown in Supplementary Fig. S2 and the quantification results of fungal hyphae are illustrated in Fig. 1. The results showed that no genomic DNA of *E. vermicola* was detected in non-infected PWNs. A small amount of fungal DNA was detected in the infected-PWNs that were under the adhesion and infection stage, as well as the early-fungal elongation stage. The fungal hyphal quantity, as expected, was significantly increased in the stages of mid-fungal elongation, late-fungal elongation, and conidium production.

**Fig. 1.**
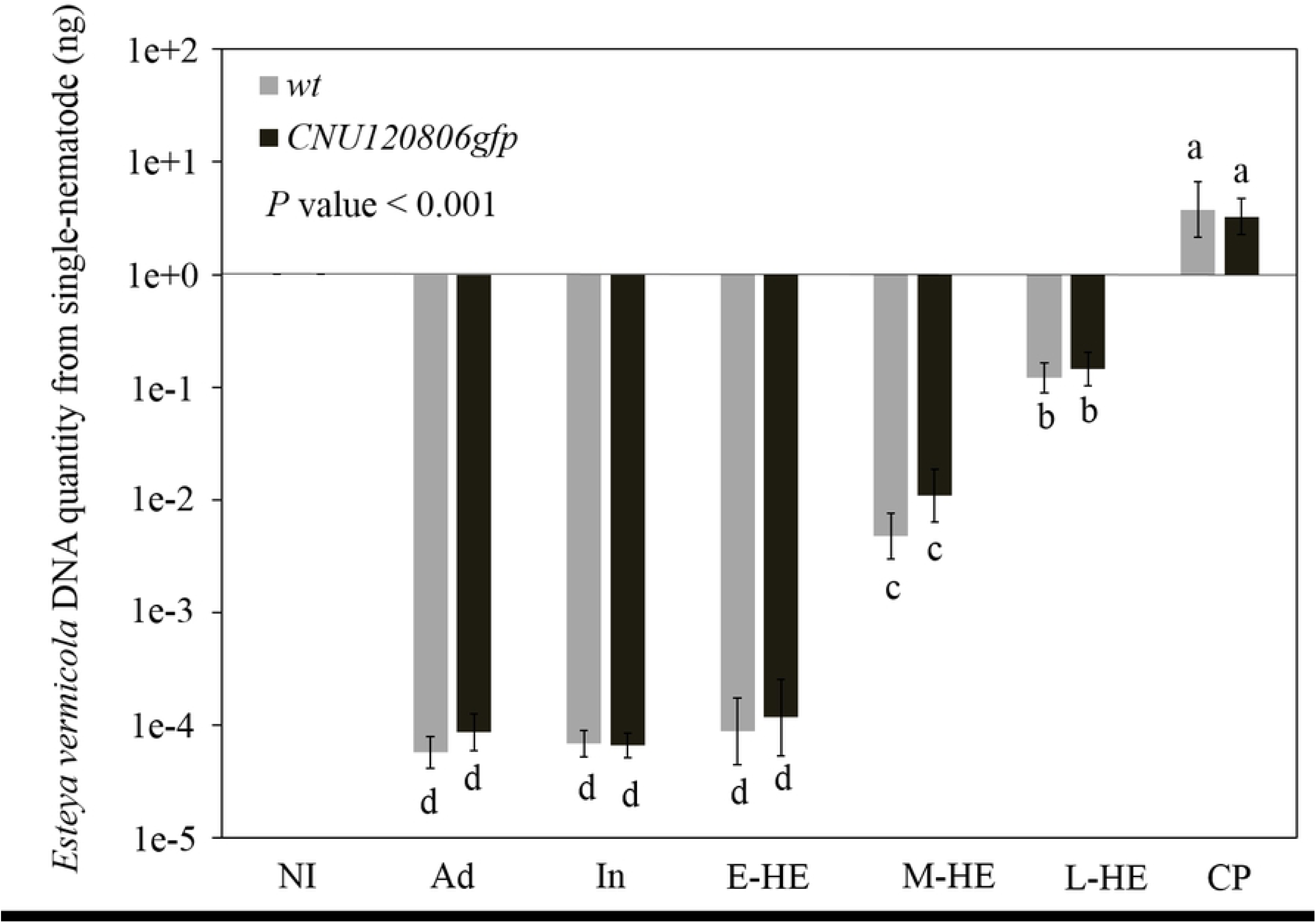
*In vitro* infection pattern of pinewood nematodes (PWNs) by *Esteya vermicola*. The values show the genomic DNA quantities in single-PWNs that were classified as various infection stages. Black columns represent the PWN infection pattern by *CNU120806gfp*; gray columns represent the PWN infection pattern by wild-type *E. vermicola* (*wt*). NI, non-infection; Ad, adhesion stage; In, infection stage; E-HE, early-hyphal elongation stage; M-HE, mid-hyphal elongation stage; L-HE, late-hyphal elongation stage; CP, conidium production stage. Columns represent the mean (± s.d.) of five biological triplicates.

#### 2.2.2 Morphological observation of parasitic fungal hyphae

The morphological characteristics of fungal hyphae parasitizing PWN were observed with the help of green fluorescence emission of GFP-tagged mutant *CNU120806gfp* (Fig. 2 and Fig. 3). According to our observations, fungal infection occurred 20 hours after the lunate conidia adhered to the cuticle of the PWN. An ovoid propagule was produced and implanted into the pseudocoelom of PWN by the fungal lunate conidia (Fig. 2a-c). The propagule found in the PWN pseudocoelom showed morphological features similar to those of fungal cells germinated by the lunate conidia cultured on a PDA plate (Fig. 2d-e). In addition, a papillary bulge was found at the center of the concave side of the germinated lunate conidia (Fig. 2d). Additionally, a similar structure was found at the germinated lunate conidia that were cultured on the PDA plate (Fig. 2e). The dumbbell-like fungal cells produced by the ovoid propagule were found in the PWN (Fig. 3a-d), which was significantly different from the fungal hyphae grown either on PDA plates (Fig. 3e) or in PDB fermentation (Fig. 3f) in morphological features. The dumbbell-like fungal hyphae released from infected PWNs were cultured *in situ* for morphological comparison. The newly produced fungal cells, as expected, showed morphological features similar to those of the fungal hyphae grown on PDA plates (Fig. 3d).

**Fig. 2.**
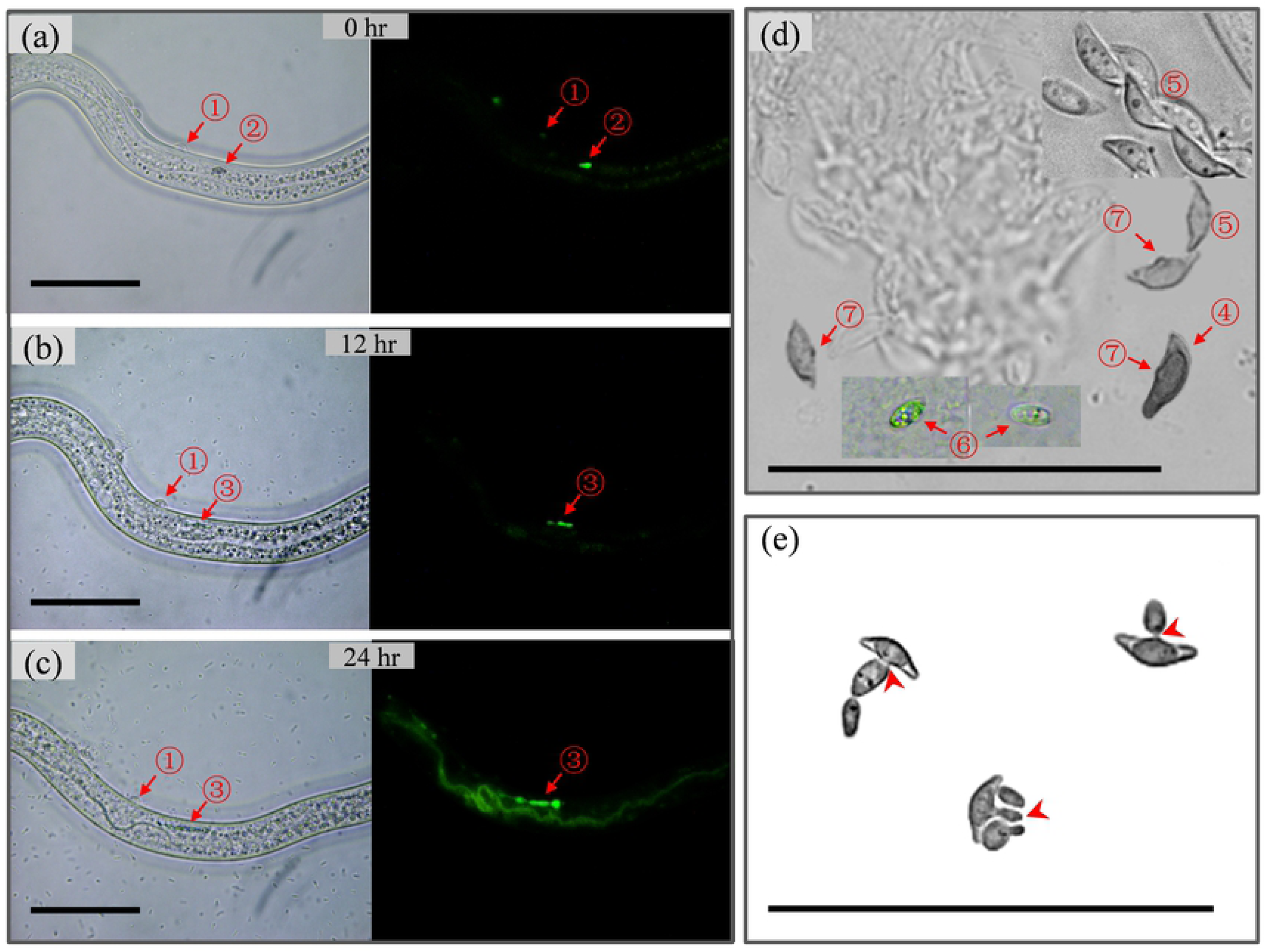
Implantation of an ovoid propagule coelom by lunate conidia triggers fungal infection of pinewood nematodes (PWNs). (a-c) The early stage of fungal infection against PWN by *Esteya vermicola*; (d) morphological characteristics of ovoid propagule and germinated lunate conidia related to the fungal infection of PWN; (e) *in vitro* germination of lunate conidia on the PDA plate. ① Lunate conidium attached on the cuticle of PWN; ② ovoid propagule implanted into the coelom of PWN; ③ growth and extension of fungal hyphae in the coelom of PWN; ④ germinated lunate conidia related to fungal infection of PWN; ⑤ ungerminated lunate conidia; ⑥ ovoid propagule released from fungal infected PWN; ⑦ papillary bulge at the center of the concave side of the germinated lunate conidia. Bar, 10 μm.

**Fig. 3.**
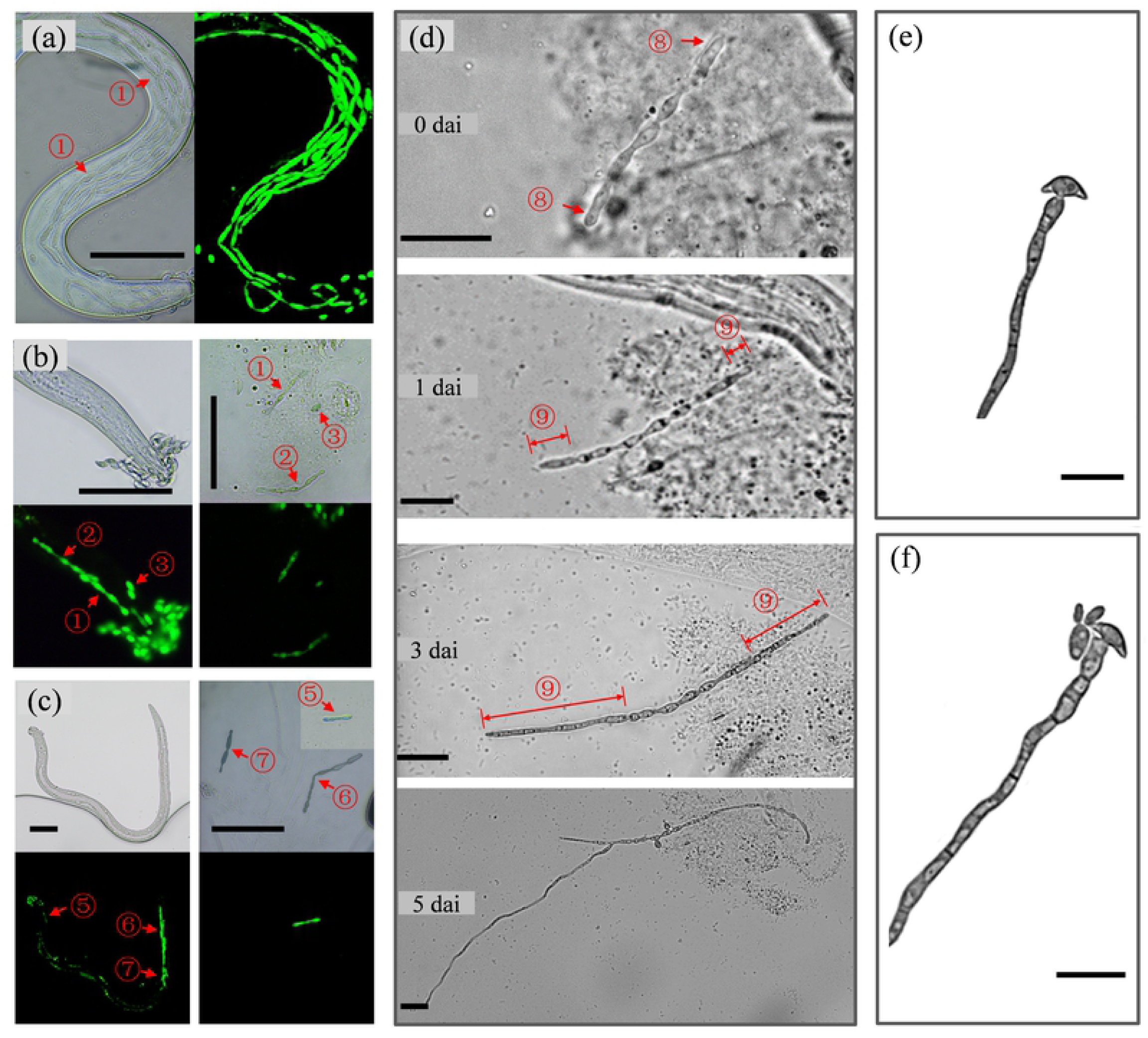
Visualization of specialized parasitic hyphae in the fungal infection of pinewood nematodes (PWNs). (a) Observation of dumbbell-like specialized parasitic hyphae in the fungally infected PWNs by *Esteya vermicola*; (b - c) the fungal hyphae in the coelom of infected PWNs; (d - e) the specialized parasitic hyphae released from the infected PWNs in (b) and (c), respectively; (f) *in situ* cultivation of the specialized parasitic hyphae released from the infected PWNs; (g) the fungal hyphae grown on PDA plates; (h) the fungal hyphae grown in PDB liquid fermentation. ① Dumbbell-like specialized parasitic hyphae in infected PWNs; ② - ⑥ dumbbell-like specialized parasitic hyphae either in or out of infected PWNs; ⑦ ovoid propagules either in or out of infected PWNs; ⑧ dumbbell-like specialized parasitic hyphae released from infected PWNs; ⑨ fungal hyphae produced by dumbbell-like specialized parasitic hyphae under *in vitro* culture conditions. Bar, 10 μm.

### 2.3 Fungal colonization of healthy susceptible pine trees by *E. vermicola*

#### 2.3.1 Temporal and spatial dynamics of *E. vermicola* colonization in healthy pine xylem

To examine the colonization patterns of *E. vermicola*, a blastospore suspension of *CNU120806gfp* was used for the inoculation of pine tree xylem. Real-time TaqMan PCR assays of DNA extracts from wood with primers and probes were conducted to discover the temporal and spatial dynamics of *E. vermicola* colonization. After inoculation, the conidia of *E. vermicola* accumulated in the wood within 1 cm of the injection site (Fig. 4a). Fifteen days later, *E. vermicola* was detected in the wood 2 cm away from the injection site (Fig. 4b). Wood samples within 5 cm of the injection site harbored this nematophagous fungus 2 months after inoculation (Fig. 4c-e). Although a small number of fungal hyphae were found in the pine tree, the extension of *E. vermicola* in the pine tree xylem at lengths indicated that the fungus germinated and grew inside the host plants. Although the relative quantities were low, as expected, the fungal hyphae were found at 90 and 180 dai, suggesting that the fungus *E. vermicola* can stably survive in host pine trees (Fig. 4e-f). Additionally, a slight increase was found at 6 months after inoculation.

**Fig. 4.**
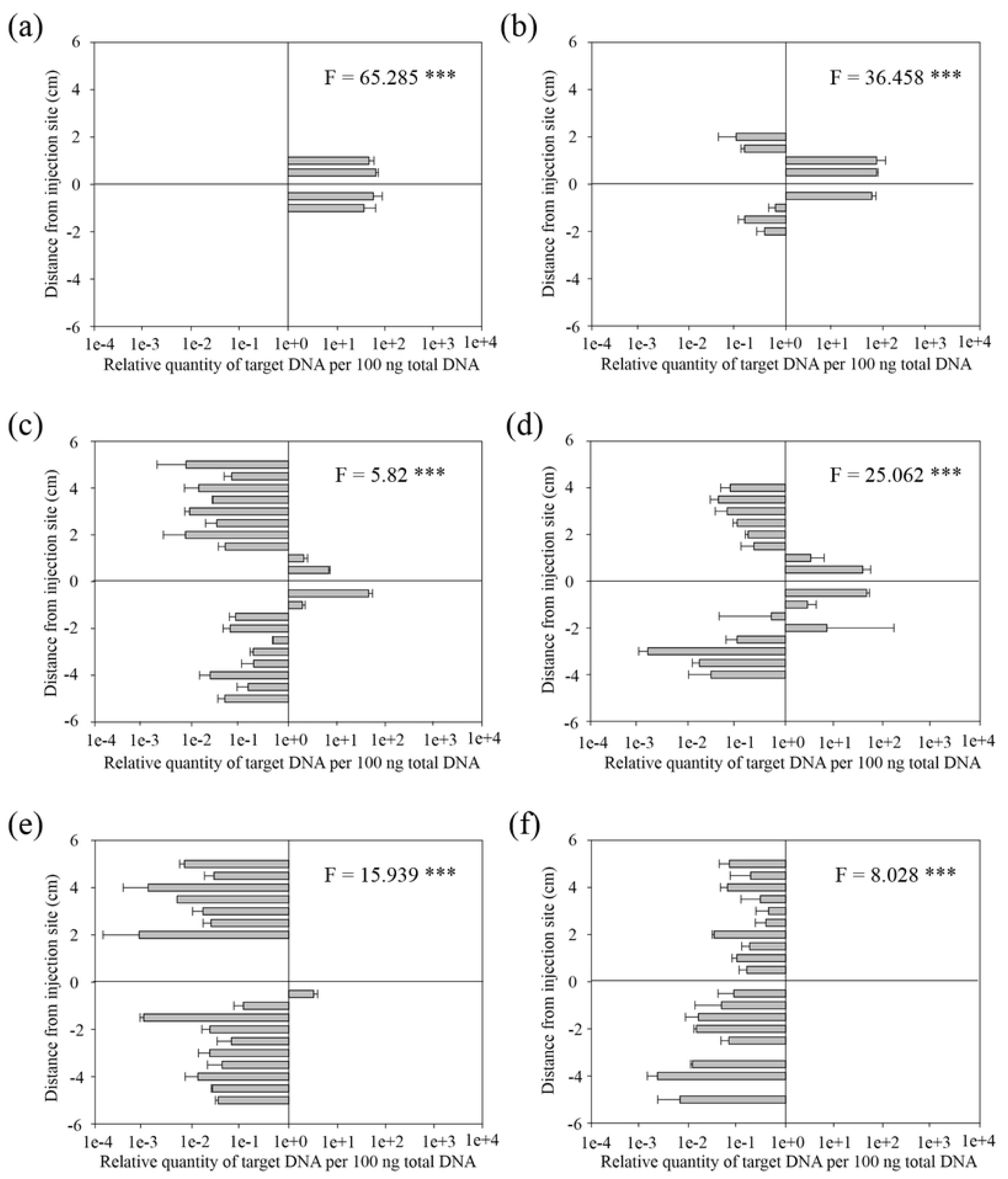
Temporal and spatial dynamics of *E. vermicola* colonization of healthy pine tree xylem. The wood samples were taken from inoculated trees within 5 cm around the inoculation site, both upward and downward. (a - f) The quantities of *E. vermicola* in pine tree xylem at 0, 15, 30, 60, 90, 180 dai, both upward and downward of the inoculation site. Y-axis, 0 represents the inoculation site, positive values represent the wood samples taken upward of the inoculation site, negative values represent the wood samples taken downward of the inoculation site. *, *P* value < 0.05; **, P value < 0.01; ***, *P* value < 0.001. Columns represent the mean (± s.d.) of three biological triplicates.

With the help of GFP fluorescence emission under a microscope, the visualization of fungal germination and colonization of *E. vermicola* is shown in Fig. 5a-b and Supplementary Fig. S3. The conidia of mutant *CNU120806gfp* were germinated after inoculation into the host tree trunk (Supplementary Fig. S3a-c), which led to a subsequent fungal hyphal colonization. Moreover, newly produced lunate conidia, which are mainly responsible for the effectiveness of fungal infection of PWNs, were found in the inoculated tree xylem (Fig. 5a).

**Fig. 5.**
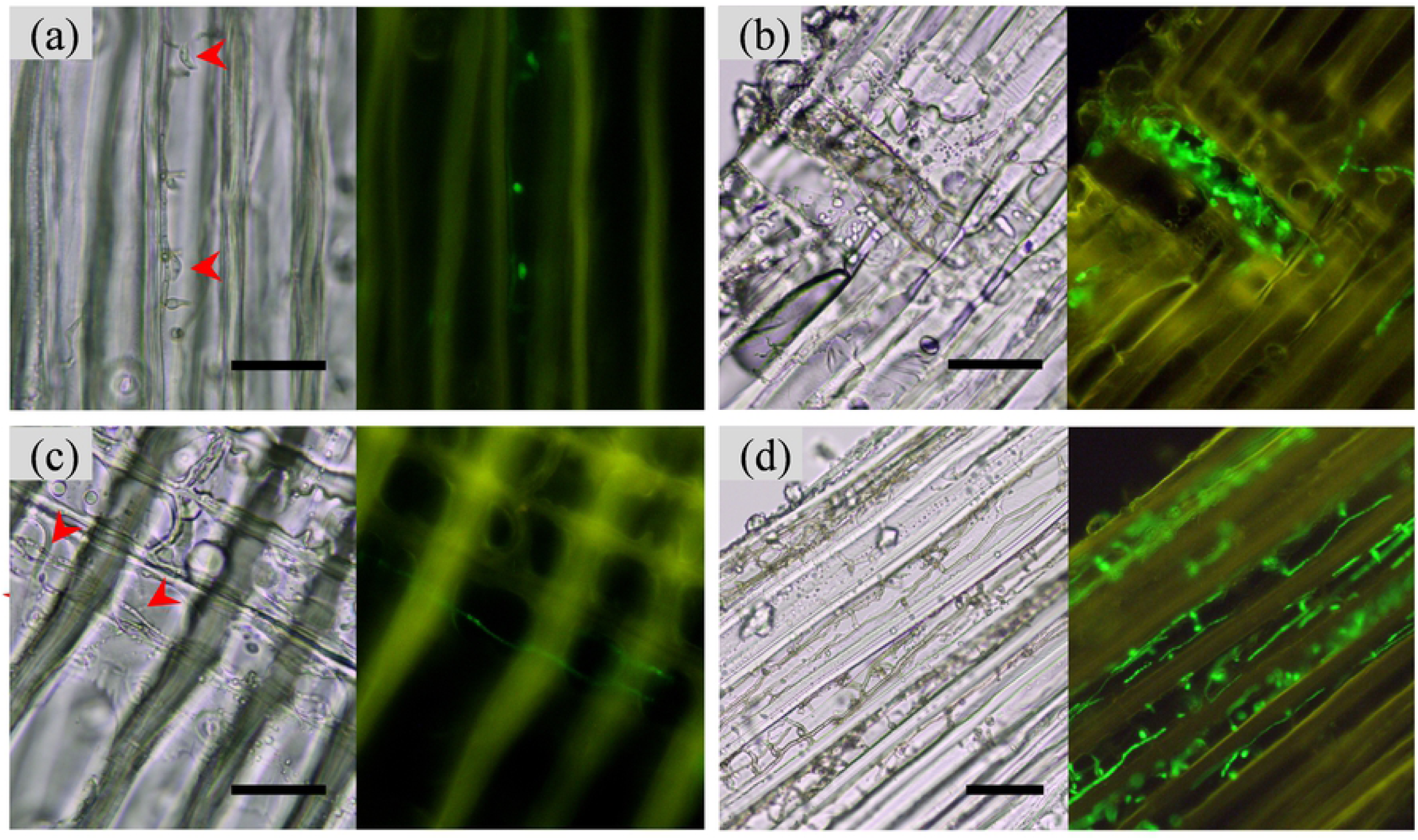
Colonization of pine tree xylem by *Esteya vermicola* with GFP. Fluorescence microscopy analyses of wood sections of pine tree xylem. (a-b) Fungal hyphae and newly produced lunate conidia of *E. vermicola* observed in healthy pine tree xylem; (c) fungal hyphae of *E. vermicola* observed in the xylem of healthy pine seedling; (d) fungal hyphae of *E. vermicola* observed in pinewood nematode-infected wilting pine tree. Bar, 20 μm.

#### 2.3.2 Temporal dynamics of *E. vermicola* colonization of pine seedling xylem and its response to PWN invasion

The xylem of pine seedlings was colonized by the *E. vermicola* mutant *CNU120806gfp* via artificial wounds on the branches. After spraying to the surface of wounds, the fungus *E. vermicola* was detected in the xylem from 7 to 28 dai (Fig. 6). According to our quantifications, however, this biological control agent cannot be detected at 45 and 60 dai. In addition, the fungal hyphae of *CNU120806gfp* were observed in wood samples from the tested pine seedlings (Fig. 5c). Although the frequency of fungal hyphae that appeared in wood sections of tested pine seedlings was very low, the occurrence of fungal hyphae suggested successful colonization by the fungus *E. vermicola*.

**Fig. 6.**
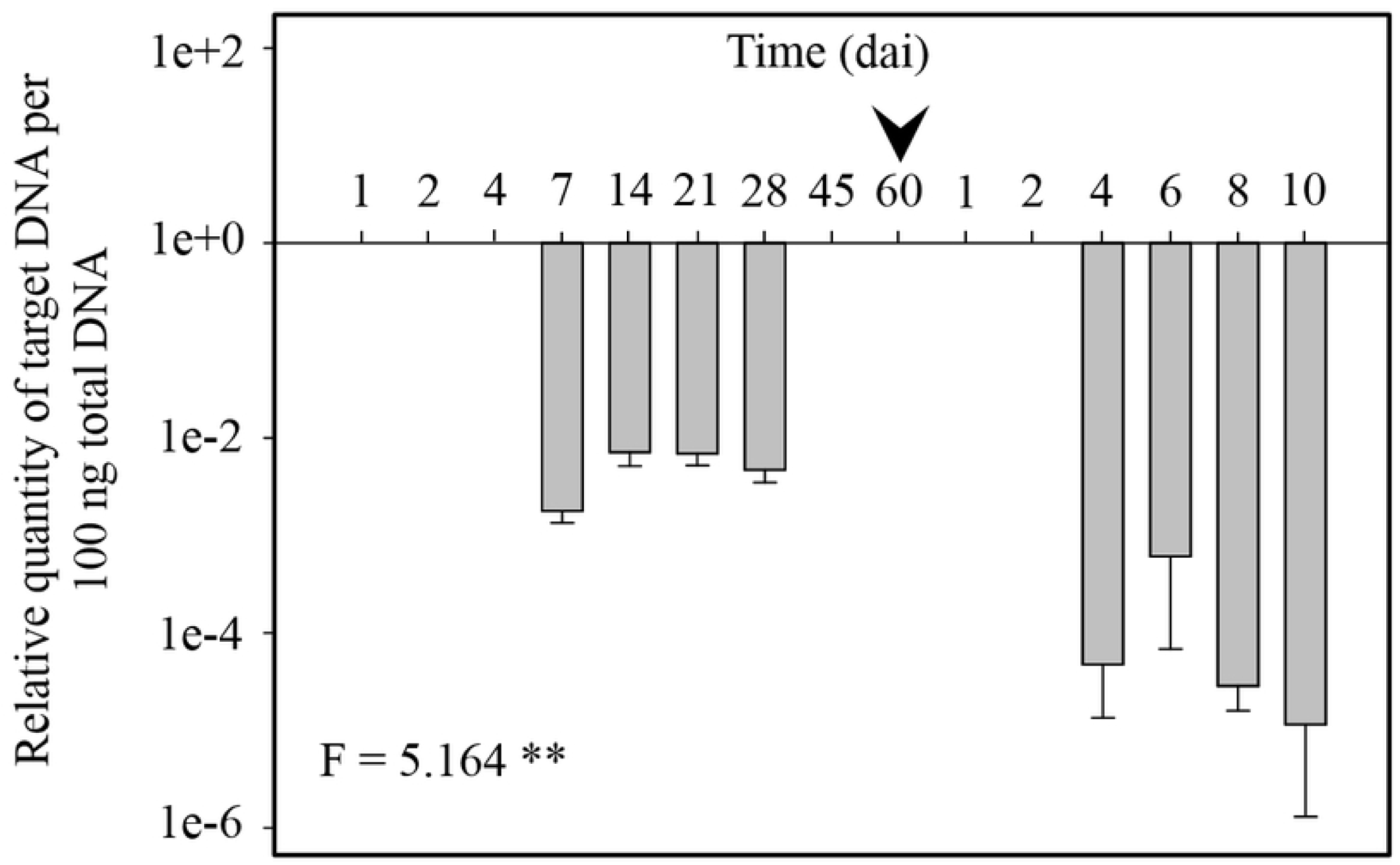
Temporal dynamics of *E. vermicola* colonization after spraying to pine seedlings and its response to pinewood nematode invasion. The quantities of *E. vermicola* in pine seedlings at 1, 2, 4, 7, 14, 21, 28, 45 and 60 days after spray and the quantity changes of *E. vermicola* in pine seedlings in response to artificial PWN infection (arrowhead) at 1, 2, 4, 6, 8 and 10 days after infection. *, *P* value < 0.05; **, *P* value < 0.01; ***, *P* value < 0.001. The column represents the mean (± s.d.) of three biological triplicates.

To reveal the response of *E. vermicola* in pine seedlings to PWN invasion, the seedlings were infected with PWNs at 60 days after fungal inoculation (Fig. 6, black arrowhead). The results showed that the quantities of *E. vermicola* slightly increased from 4 days after PWN infection, although in extremely low quantities. This result suggested that *E. vermicola* inhabited the pine tree xylem as a low-richness species; however, *E. vermicola* quantities increased when PWN invasion occurred.

The PWNs were extracted from wood samples of these tested seedlings. *B. xylophilus* was first extracted from pine seedlings at 2 dai, including the fungus-infected and uninfected PWN (Fig. 7a and 7b). In the pine seedlings inoculated with *E. vermicola*, the total number of extracted PWNs increased within 6 dai and then decreased from 8 dai onward (Fig. 7a); while in the control group, the number of extracted PWNs increased continuously after inoculation (*P* < 0.01) (Fig. 7b). The PWN infected by *CNU120806gfp* extracted from pine seedlings was visualized using white light and fluorescence microscopy (Fig. S4a). In most cases, the infected PWN was in the early stage of fungal infection by *E. vermicola*. The relative quantity of *E. vermicola* assayed by TaqMan probe quantification illustrated the infection status of extracted PWN from pine seedlings (Fig. 7c). The relative quantity of *E. vermicola* DNA in PWNs recovered from fungus-treated plants increased from 4 dai, and reached the top value at 10 dai. Subsequently, the relative quantity of fungal genomic DNA was decreased followed by a decline in the PWN amount recovered from pine seedlings from 12 dai.

**Fig. 7.**
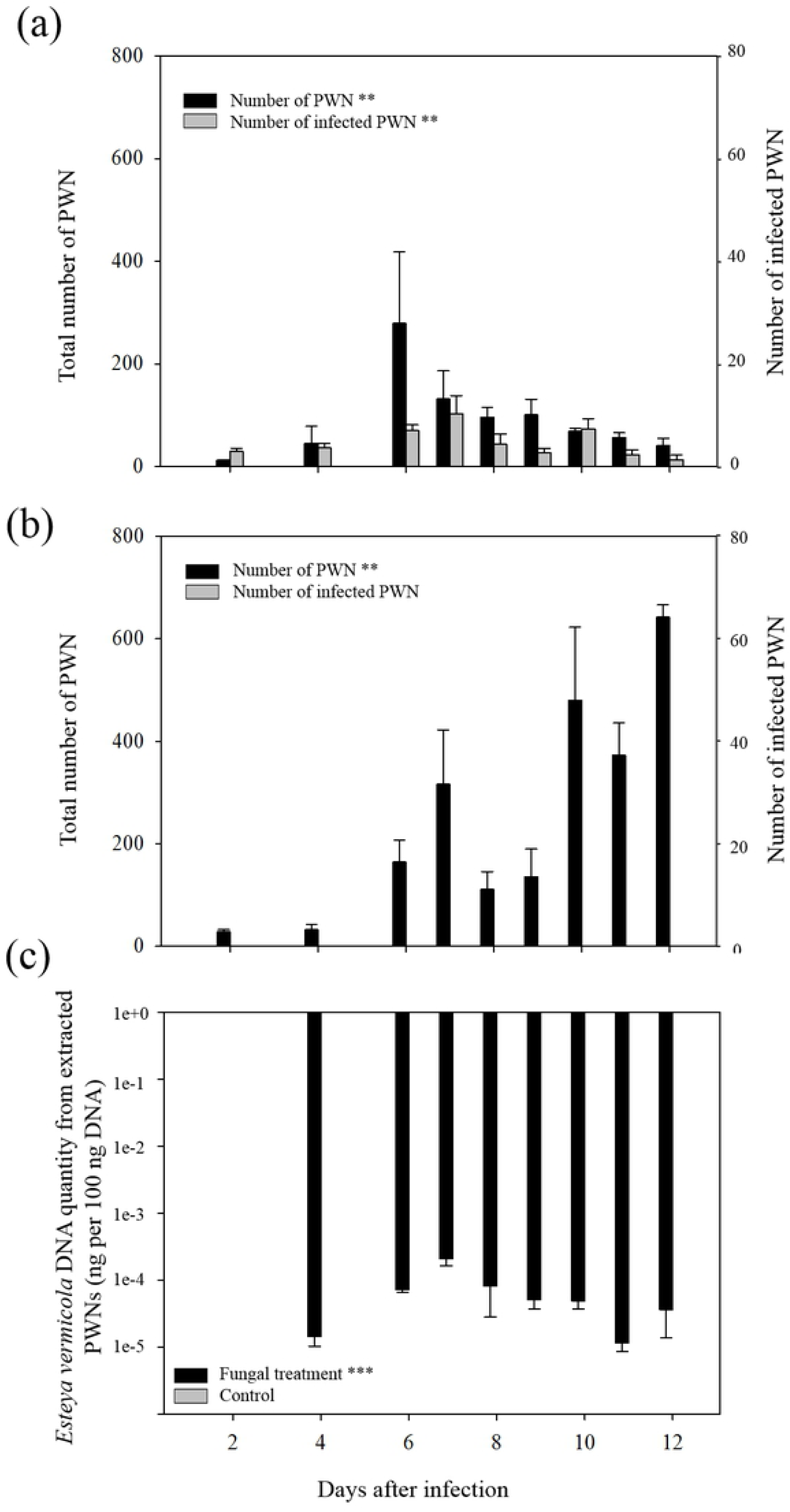
Fungal infection status of pinewood nematodes (PWNs) extracted from pine seedlings. (a) The number of infected and non-infected PWN extracted from *Esteya vermicola* inoculated pine seedlings; (b) the number of PWN extracted from control pine seedlings. Black column, the total number of PWN extracted from pine seedlings; gray column, the number of fungal infected PWN extracted from pine seedlings. (c) The relative quantity of fungal genomic DNA of *E. vermicola* in PWNs extracted from pine seedlings. Black column, the relative fungal quantity of *E. vermicola* in PWN extracted from fungal inoculated pine seedlings; gray column, the relative fungal quantity of *E. vermicola* in PWN extracted from control pine seedlings. *, *P* value < 0.05; **, *P* value < 0.01; ***, *P* value < 0.001.

### 2.4 Colonization and infection patterns of *E. vermicola* in the wilting pine tree

#### 2.4.1 Temporal and spatial dynamics of *E. vermicola* colonization in wilting pine xylem

The quantification of the target gene in wood samples revealed the colonization patterns of the fungus *E. vermicola* in PWN-infected wilting pine trees (Table 1). The results indicated that *E. vermicola* was only detected in the wood tissue around the inoculation site within 7 dai. Two weeks later, however, the fungus *E. vermicola* was detected 10 cm away from the inoculation point. Subsequently, the genomic DNA of *E. vermicola* was found in the xylem 50 cm away from the inoculation point at 21 dai. However, the fungal colonization patterns showed jumping spread in the wilting pine tree. The extension patterns that appeared in wilting plants were thought to be due to the immigration of parasitized PWNs in the pine xylem. The hyphal morphology of *E. vermicola* inoculated in PWN-infected wilting pine xylem was observed (Fig. 5d). The fungal hyphae in the wilting pine tree showed higher abundance and more irregular development than those in healthy trees.

**Table 1.**
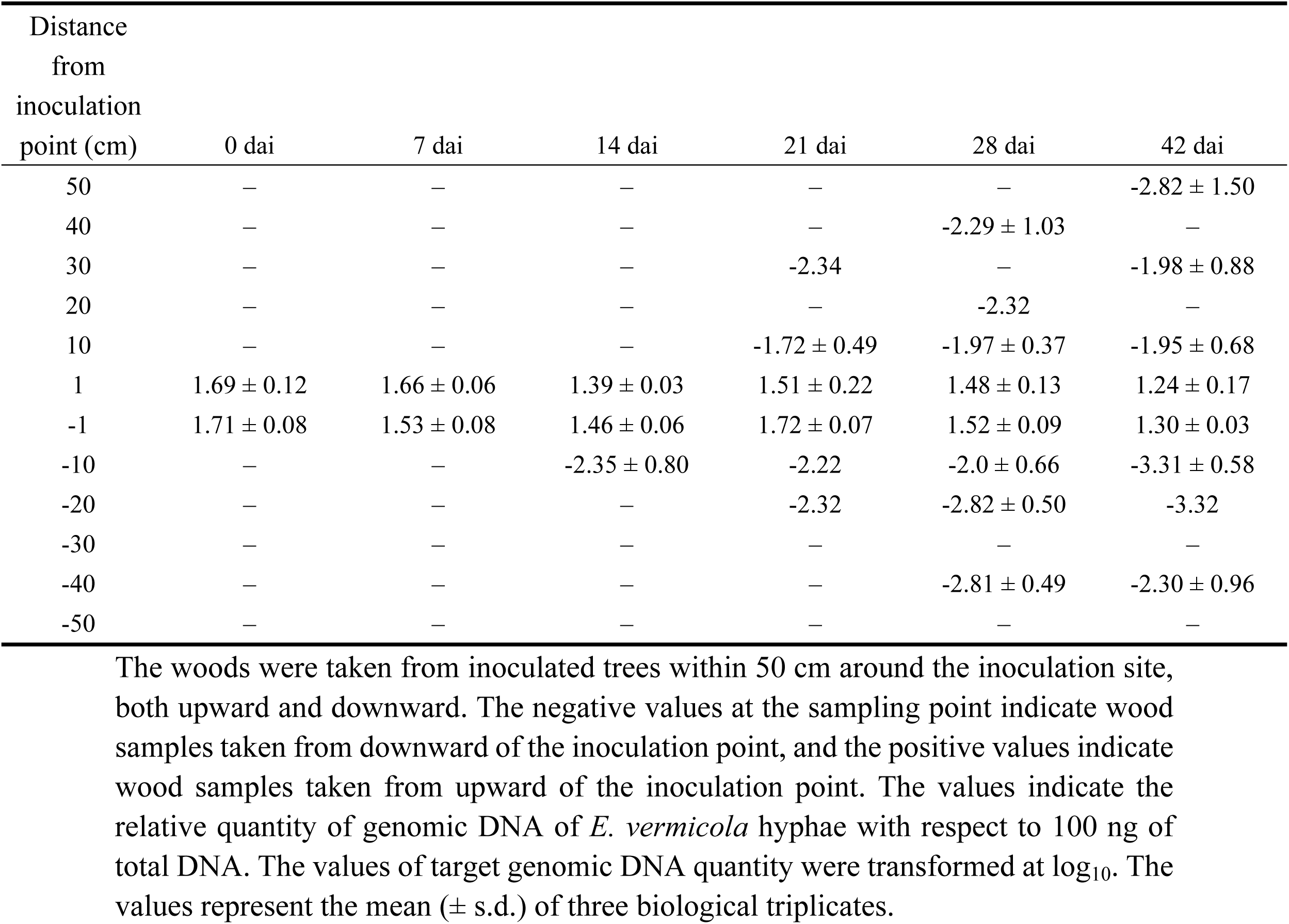
Temporal and spatial dynamics of *Esteya vermicola* colonization of pinewood nematode-infected wilting pine trees.

#### 2.4.2 PWN density in the wilting pine tree

The PWNs inhabiting wilting host plants were measured in the present study, and the results are shown in Fig. 8. As shown in the figures, the PWN density in the xylem of both *E. vermicola* and sterile ddH_2_O-treated wilting pines was increased. However, the number of PWNs per 10 g of the wood sample around the *E. vermicola* inoculation point decreased from 7 dai to 21 dai (Fig. 8a-d), and PWN could not be extracted at 28 and 42 dai (Fig. 8e-f). Additionally, the PWN density in the wood 10 and 20 cm away from the inoculation site significantly declined after fungal inoculation (Fig. 8c-f).

**Fig. 8.**
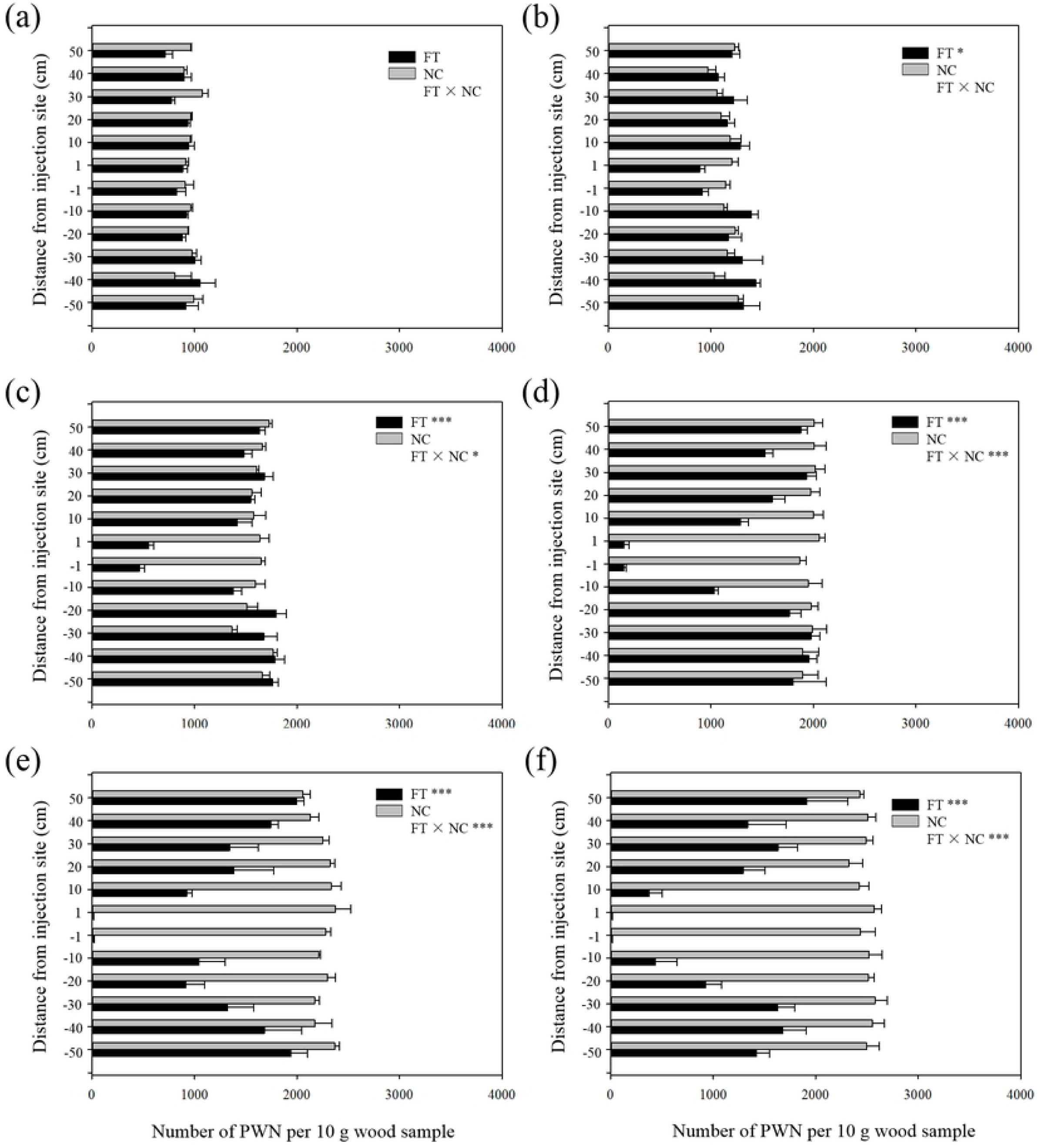
Relative quantity of fungus *E. vermicola* in the pinewood nematode (PWN) isolated from wilting pine woods. The wood samples were taken from inoculated wilting trees within 50 cm around the inoculation site, both upward and downward. (a - f) The quantities of *E. vermicola* in pinewood nematodes isolated from the plants at 0, 7, 14, 21, 28, and 42 dai, both upward and downward of the inoculation site. FT, fungal treatment; NC, negative control. *, *P* value < 0.05; **, *P* value < 0.01; ***, *P* value < 0.001. Columns represent the mean (± s.d.) of three biological triplicates.

#### 2.4.3 Fungal quantity of *E. vermicola* in extracted PWNs

Additionally, the amount of *E. vermicola* in the body of extracted PWNs was quantified using TaqMan PCR quantification (Table 2). The results showed that *E. vermicola* could be detected in PWNs extracted from wood samples taken from 7 dai. The fungal hyphae, as expected, could not be detected in the PWN extracted from the wilting trees treated with sterile ddH_2_O. Similar to the extension of *E. vermicola* in wilting pine trees, the presence of fungal hyphae in extracted PWNs showed jumping spread. These results suggested that the fungal colonization and extension of *E. vermicola* in PWN-infected pine trees were significantly correlated with PWN immigration in the xylem. This might result in accelerated fungal diffusion in the host plant and increased fungal infection of PWNs located throughout the tested tree. To illustrate the fungal infection of PWN extracted from the wilting pine tree, the hyphae of *E. vermicola* were visualized under a fluorescence microscope (Supplementary Fig. S4b).

**Table 2.** The relative quantity of *Esteya vermicola* in pinewood nematodes extracted from fungal-inoculated wilting pine trees.

| Distance from inoculation point (cm) | 0dai | 7dai | 14dai | 21dai | 28dai | 42dai |
| --- | --- | --- | --- | --- | --- | --- |
| -50 | — | — | — | — | — | -5.65 ± 0.70 |
| -40 | — | — | — | -4.16 ± 0.17 | -3.92 ± 0.07 | -3.32 |
| -30 | — | — | — | — | -3.37 | -4.57 ± 0.41 |
| -20 | — | — | -3.82 ± 0.50 | -2.32 | -3.84 ± 0.82 | -3.35 |
| -10 | — | -4.32 | -3.32 ± 0.58 | -3.87 ± 0.30 | -2.97 ± 0.62 | -4.66 ± 0.34 |
| -1 | — | -2.82 ± 0.50 | — | — | — | — |
| 1 | — | -2.84 ± 1.48 | — | — | — | — |
| 10 | — | — | — | -3.66 ± 0.32 | -3.86 ± 0.35 | -2.99 ± 0.66 |
| 20 | — | — | -4.32 | -4.32 | -3.40 ± 0.25 | — |
| 30 | — | — | -4.32 | -3.84 ± 0.49 | -4.27 ± 0.20 | -3.51 ± 0.24 |
| 40 | — | — | — | -3.67 ± 0.35 | -4.15 | -3.28 ± 0.38 |
| 50 | — | — | — | — | -4.18 ± 0.82 | -4.10 ± 0.44 |

The pinewood nematode was extracted from the wood of fungal inoculated wilting trees within 50 cm around the inoculation site, both upward and downward. The negative values at the sampling point indicate wood samples taken from downward of the inoculation point, and the positive values indicate wood samples taken from upward of the inoculation point. The values indicate the relative quantity of genomic DNA of *E. vermicola* hyphae with respect to 100 ng of total DNA. The values of target genomic DNA quantity were transformed at $\log_{10}$ . The values represent the mean ( ± s.d.) of three biological triplicates.

## 3 Discussion

Fungal colonization and *in vivo* infection of *B. xylophilus* by *E. vermicola* inside susceptible pine trees is crucial for the biocontrol of PWD. Moreover, fungal colonization of the susceptible pine xylem before potential PWN invasion can significantly improve the control effects [8,9]. For a biological agent that infects and controls PWN in pine trees, the colonization and extension of *E. vermicola* inside of its host tree are essential. However, the temporal and spatial dynamics of *E. vermicola* colonization of the host pine xylem are unknown in both healthy and wilting pine trees. Thus, a time-course investigation on the presence of *E. vermicola* in healthy and PWN-infected pine trees is important to understand fungal colonization and is helpful to improve forest management.

By visualizing the parasitic hyphae in infected PWNs, the specific infection approach used by *E. vermicola* to infect *B. xylophilus* was explored. Although the infection of *B. xylophilus* by *E. vermicola* has been studied [11,19], the specific implantation method and specialized fungal hyphae in PWN infection were first reported in the present study. The infection process of the plant-parasitic nematode by endoparasitic fungi has been investigated using the GFP-labeled fungal transformant [13,20,21]. The fungal infection of these plant-parasitic nematodes began with the formation of an infection peg, and then the nematode coelom was colonized by fungal hyphae. Different from the modes used by most nematophagous fungi to infect host nematodes, the fungal infection of *B. xylophilus* by *E. vermicola*, however, began with the implantation of an ovoid propagule by the germinated lunate conidia that adhered to the cuticle of PWN. Subsequently, dumbbell-like fungal cells were produced and grown through the coelom of PWN, which is significantly different from the fungal hyphae grown in either PDA or fermentation cultures. The morphological features of hyphal growth out of the coelom and *in situ* culture of the released dumbbell-like fungal hyphae suggest that specialized fungal cells were produced only when parasitizing PWNs. However, the functions of specialized fungal cells in infecting *B. xylophilus* remain unknown. The temporal infection dynamic of PWN by *E. vermicola* was explored with TaqMan PCR quantification. No significant increase in fungal hyphal quantity was found before the early stage of hyphal elongation. However, the fungal abundance was significantly increased from mid-hyphal elongation, suggesting the rapid growth of fungal hyphae by degrading the organs and tissues of PWNs.

Spatial and temporal analyses of *E. vermicola* within healthy pine tree xylem using TaqMan qPCR techniques revealed the patterns and dynamics of fungal colonization and the extension rate inside pine trees. After inoculation, the fungal conidia accumulated within 1 cm of the inoculation sites. This result validates the hypothesis that the fungal conidia of *E. vermicola* cannot pass through the pits on the tracheid wall of a healthy pine tree [11]. Subsequently, the conidia of *E. vermicola* were germinated and grew within a few days after inoculation and extended upward and downward. A small quantity of fungus, however, extended to a sampled site more than 1 cm away from inoculation sites. Although the extension rate is slow, fungal hyphae of *E. vermicola* were indeed growing through the wall of the tracheid. Regarding the low quantity of fungal hyphae detected in the farther samples from inoculation sites, the inhibitory effects of plant defense compounds and segregation of the tracheid wall [22,23] are supposed to be the main limiting factors. Moreover, the fungal hyphae were detectable and observable at 180 dai after fungal inoculation, suggesting that the fungal hyphae could stablely colonize pine tree xylem.

The fungal colonization of pine seedlings by *E. vermicola* was investigated, and the pine seedling stem can be colonized via artificial wounds [9]. After spraying inoculation of the wounds, as expected, the quantity of *E. vermicola* inhabiting the pine seedling xylem showed a moderate increase. However, the target DNA could not be detected at 45 and 60 dai. The colonization behavior detected for *E. vermicola* was probably responsible for the inhibition of the plant defense systems, as described in previous studies [24,25]. Then, the inoculated pine seedlings were infected with PWN suspension. The quantity of *E. vermicola* increased in response to PWN invasion in our study, indicating that the fungal hyphae in pine tree xylem parasitized the invading PWNs and led to a moderate increase in fungal quantity from the undetectable value. Previous reports suggested that invasion of PWNs would enhance the defense system of the host plant due to the destruction of plant tissues [22,26]. However, the fungal quantities were moderately increased after PWN invasion in our study, despite increasing defense compound levels in the pine seedlings. The increase in fungal quantity could have occurred due to the elicitation of PWN infection cycles by *E. vermicola* once the PWNs invaded and were exposed to lunate conidia, providing nutrients for fungal propagation. These results are supported by the observation of fungal hyphae and successful extraction of fungal-infected PWNs.

Regarding the inoculation of wilting pine trees, the fungal hyphae of *E. vermicola* extended rapidly and illustrated a jumping spread in the pine xylem. Similar to the fungal inoculation of healthy pine trees, the conidia inoculated with wilting pine xylem also accumulated around the inoculation sites, followed by rapid germination and colonization. However, the PWNs in the wilting pine trees supply plenty of nutrients for the growth of the nematophagous fungus *E. vermicola*. Thus, the fungal hyphae were extended rapidly. Moreover, PWNs parasitized by *E. vermicola* do not die within 3 days [27]. The PWNs attached by lunate conidia or infected by *E. vermicola* can move in the first 2 days after infection. In this case, the jumping spread of *E. vermicola* is suggested to be triggered by the immigration of infected PWNs, which was confirmed by the distribution pattern of fungi detected in wood samples, as well in extracted PWNs. Furthermore, the significant decline in PWN density in wilting pine xylem after inoculation with *E. vermicola* indicates the great potential of this biological control agent in the management of PWD.

It is well known that resin is the main chemical defense substrate in conifers and that it includes the volatile compounds α-pinene, β-pinene, myrcene, limonene, and 3-carene [28,29], which injure or kill beetles, nematodes and other invaders. According to a previous study, these compounds are the main volatile monoterpenes obtained from resin [30], and the amount of these compounds increases after an insect or artificial attack [31]. Likewise, the growth of fungi is inhibited by the volatile compounds in the resin [32]. A comparative genomic study indicated that the gene family expansion of xenobiotic detoxification pathways, including flavin monooxygenase (FMO), Cytochrome P450 (CYP450), short-chain dehydrogenase (SDR), alcohol dehydrogenase (ADH), aldehyde dehydrogenase (ALDH), UDP-glucuronosyltransferase (UGT) and glutathione S transferase (GST), facilitated the survival of PWN in pine resin ducts [33,34]. Additionally, the orthologous genes related to aspartic endopeptidase, aminopeptidase, and CYP450 reductase revealed positive selection in *E. vermicola* compared with insect pathogenic and plant pathogenic fungi [35]. The moderate expansion of detoxification-related gene families is supposed to be a crucial feature as a biocontrol agent against *B. xylophilus*. However, the molecular detoxification mechanisms of the *E. vermicola* response to pine defense systems are less known, and investigations are required in future studies.

To the best of our knowledge, we are the first to investigate the colonization patterns of the endoparasitic fungus *E. vermicola* in host pine tree xylem. Simultaneously, the specific infection method used by *E. vermicola* infects *B. xylophilus*, and specialized fungal parasitic cells in PWN infection are first reported. The study revealed the spatial and temporal colonization dynamics of this plant nonpathogenic fungus in healthy and PWN-infected wilting pine trees, as well as its responses to PWN invasion. The results provide informative knowledge about the extension rate of this fungus in the biological control of PWN and will be helpful for the management of PWN in the field. Although the high infectivity against PWN and effectiveness in controlling PWD by *E. vermicola*, fungal colonization is supposed to need a longer time to extend throughout the susceptible trees. This means that a longer colonization time is needed after fungal inoculation to achieve the highest biocontrol effects against PWN invasion. Moreover, the development of rapid colonization techniques is essential by using either innovation of the inoculation method or genetic modification of the *E. vermicola* genome.

## 4 Materials and methods

### 4.1 Fungal strain, nematodes, and host plants

The experiments were conducted with an isolate of *E. vermicola*, CNU120806, isolated from infected PWNs in South Korea. In addition, the mutant *CNU120806gfp* with the *gfp* gene was used to visualize and locate the fungal hyphae in the tested plants. The characteristics of the GFP transformant strain *CNU120806gfp*, including growth rate, conidial yield in fermentation culture, conidial germination rate, and infection rate of PWN, are similar to those of the wild strain [11]. *E. vermicola* strains were grown in potato dextrose agar (PDA; KisanBio Co., Seoul, Korea) at 26 °C in an incubator for 10 days, or in potato dextrose broth (PDB; KisanBio Co., Seoul, Korea) at 26 °C, and 120 rpm for 7 days on a shaker. The fungal blastospore suspension used in the experiments was harvested from 7-day culture fermentation of either the wild type (*wt*) strain or mutant *CNU120806gfp*. Then, the blastospore suspension was washed three times using sterile distilled water by centrifugation to remove the residual nutrient content.

The *B. xylophilus* used in this study was isolated from a wild PWN-infected forest on Jeju Island, South Korea, and cultured on fully grown *Botrytis cinerea* plates. The PWNs were extracted from plates using the Baermann funnel technique [36] after the fungus *B. cinerea* was completely eaten. Then, the PWN suspensions were clean with sterile distilled water three times to remove fungal fragments before use in subsequent experiments. Additionally, 10-year-old and 4-year-old *Pinus koraiensis* grown in a greenhouse were used in this study.

### 4.2 Real-time PCR quantification and standard curves

To quantify the fungal hyphae of *E. vermicola* in the tested samples, the TaqMan probe PCR quantification technique combined with nested-based PCR developed by Wang et al. (2020a) [18] was used. A specific fragment on the 28S region of the ribosomal RNA was used for the design of the primers and probes. The primers 28S-1F/28S-1R were used in the first-round PCR to amplify a 473 bp fragment. Based on the amplified fragment, the sequences of primers 28S-2F/28S-2R and probe Ev-28SP were used for second-round TaqMan PCR quantification. For the TaqMan probe qPCR, amplifications were performed in a total volume of 20 μl using an ExicyclerTM 96 Real-Time Quantitative PCR System (Bioneer, Daejeon, South Korea). The amplification results were analyzed using ExicyclerTM 96 Detection and Analysis software (version 3.0). The primers and probe are listed in Supplementary Table S1.

In the present study, two calibration curves targeting different concentration ranges of target DNA were constructed to quantify both *wt E. vermicola* and *CNU120806gfp*. Traditional TaqMan PCR was performed for the target DNA at concentrations ranging from 10^−2^ to 100 ng of total template DNA with the specific primers 28S-2F/28S-2R. In addition, nested TaqMan PCR-based quantification was performed for target DNA ranging from 10^−5^ ng to 10^−1^ ng with the primer pairs 28S-1F/28S-1R and 28S-2F/28S-2R to detect the target DNA at an extremely low concentration. The *C*_*t*_ values obtained from the reactions were correlated with the amount of DNA and were used to construct calibration curves. Three replicates were conducted for each treatment.

### 4.3 *In vitro* infection patterns of PWN by *E. vermicola*

#### 4.3.1 *In vitro* infection and fungal-infected PWN preparation

The infection of PWN by the fungus *E. vermicola* was studied under experimental conditions. The blastospore suspension (1 × 10^6^ ml^-1^) of mutant *CNU120806gfp* and *wt E. vermicola* were used for *in vitro* infection of PWNs. Herein, approximately 10^5^ fungal blastospores were spread on the water agar (WA) plates and cultured at 26 °C in an incubator for 5 days, when a large number of lunate conidia were produced by the germinated fungal hyphae. Then, approximately 1,000 PWNs were introduced to each plate with numerous lunate conidia.

The PWNs were observed and picked with a metal filament at 1, 12, 24, 36, 48, and 72 hours after PWN was introduced under white light and fluorescence microscopy (OLYMPUS, CX43, Germany). Meanwhile, the infection stage of these picked PWNs was classified as adhesion, infection, hyphal elongation, or conidium production according to the description by Wang et al.(2020b) [11]. A total of seven infection time-points of PWN (non-infection, adhesion, infection, early-hyphal elongation, mid-hyphal elongation, late-hyphal elongation, and conidium production) infected by *E. vermicola* were selected, and five PWNs were picked at each time-point. Then, the classified PWN was placed in a 1.5-ml tube with 10 μl of worm lysis buffer and preserved at −20 °C for subsequent fungal quantification.

#### 4.3.2 Quantification of *E. vermicola* in single-PWN

Total DNA was extracted from single PWN using worm lysis buffer (WLB) following the method described by Adam et al. (2007) [37] with some modification. The WLB was contained: 50 mM KCl, 10 mM Tris pH 8.2, 2.5 mM MgCl_2_, 0.45% NP40 (Fisher Scientific, Korea), 0.45% Tween 20 (Sigma, USA), 0.01% gelatin and 30 μg ml^−1^ lyticase. The PWN was placed in a 1.5-ml tube with 10 μl of WLB without proteinase K and gently ground using a grinding rod. Then, the mixture was incubated at 37 °C for 15 min and transferred by pipette into a 0.2-ml tube, and 120 μg ml^−1^ proteinase K (Bioneer, Daejeon, South Korea) was added to 15 μl. Subsequently, the tube was placed in liquid nitrogen for 5 min, and then thawed at 60 °C for 10 min. The same freeze-thaw procedure was repeated twice, followed by 90 °C for 10 min. Finally, 15 μl of DNA extract was obtained for each sample. For fungal quantification, 5 μl of total DNA extract was added to each reaction as a DNA template. The *C*_*t*_ values of *E. vermicola* quantification were obtained from either traditional or nested-based TaqMan probe PCR. Then, the *C*_*t*_ values were interpolated to the corresponding calibration curve to calculate the genomic DNA quantity of *E. vermicola*.

#### 4.3.3 Visualization of *E. vermicola* in infected PWNs

The PWNs infected by lunate conidia of *CNU120806gfp* were picked with a metal filament. Immediately, the nematodes were mounted on glass slides. The PWNs infected by the fungal hyphae were observed under the white light and fluorescence microscopy. Then, the parasitism hyphae were released from the pseudocoelom of infected PWN by physical squeeze on the cover glass. The fungal hyphae and germinated lunate conidia participating in PWN infection were visualized. Subsequently, the fungal hyphae released from the pseudocoelom of infected PWNs were cultured *in situ* at 25 °C to compare the morphological differences between fungal hyphae grown *in vitro* and *in vivo*. Simultaneously, the fungal hyphae grown on PDA plates and in PDB fermentation were observed.

### 4.4 Wood sample preparation for fungal colonization investigation

#### 4.4.1 Temporal and spatial dynamics of fungal colonization in healthy pine trees

To investigate the temporal and spatial dynamics of *E. vermicola* colonization in healthy pine tree xylem, 10-year-old *Pinus koraiensis* (approximately 10 cm in diameter) grown in a greenhouse were used for inoculation of *E. vermicola*. A total of 15 ml of blastospore suspension (3 × 10^8^ ml^-1^) of *E. vermicola* mutant *CNU120806gfp* harvested from 7-day-old liquid fermentation culture was injected into holes drilled in the trunk, and then the holes were plugged with tapered wooden stoppers. The tested trees that were inoculated with the fungal conidia suspension were cut at 0, 15, 30, 60, 90, and 180 days after inoculation (dai). Pine xylem within 5 cm of the inoculation site was sampled for the investigation of *E. vermicola* colonization. The wood samples were cut lengthways into several pieces at intervals of 0.5 cm from the inoculation hole, in both the upward and downward directions. The sampled woods were separated into two parts, one was stored at −20 °C for total DNA extraction and TaqMan PCR quantification of *E. vermicola*; another was used for the free-hand section to observe fungal colonization using a fluorescence microscope.

#### 4.4.2 Temporal dynamics of fungal colonization inside healthy pine seedlings

To study the temporal dynamics of *E. vermicola* colonization in pine tree xylem and its response to PWN invasion, a series of experiments were performed. Twenty-seven 4-year-old *P. koraiensis* pine seedlings grown in a greenhouse were used to study the temporal dynamics of *E. vermicola* colonization in pine seedlings. For fungal inoculation, the bark of pine seedling stems was randomly cut to create artificial wounds, and then a blastospore suspension (3 × 10^8^ ml^-1^) of *E. vermicola* mutant *CNU120806gfp* was sprayed onto the wounds of the seedlings. Subsequently, the wounded branch of seedlings was individually covered by cling film for 7 days to maintain moisture. After that, the pine seedlings were sampled at 1, 2, 4, 7, 14, 21, 28, 48, and 60 dai. The wood samples were used to determine the population dynamics of *E. vermicola* inside pine seedlings after inoculation, as well as to reveal the fungal location in the pine seedlings.

#### 4.4.3 Response of the fungal quantity inside pine seedlings to PWN invasion

In this experiment, eighteen 4-year-old *P. koraiensis* seedlings were treated with a blastospore suspension (3 × 10^8^ ml^-1^) of the *E. vermicola* mutant *CNU120806gfp* following the method described in the previous section and cultured in a greenhouse for two months. Subsequently, the pine seedlings were artificially infected by PWNs. Two pieces of bark (0.4 × 0.4 cm^2^) were removed from the seedling stems at 10 cm intervals, approximately 50 μl of a suspension (containing approximately 10,000 nematodes) was immediately dropped onto each of the wounds, and the wounds of the pine seedlings were covered with cling film. Simultaneously, 18 untreated pine seedlings were artificially infected with PWN suspension using the same method as the control group. The pine seedlings were sampled at 1, 2, 4, 6, 8, and 10 days after artificial PWN infection. The sampled seedling woods were used for the estimation of the fungal quality of *E. vermicola* using the TaqMan PCR quantification technique. The woods were also used for fungal visualization by fluorescence microscopy and PWN extraction with the Baermann funnel technique. The fungal infection status of the extracted PWNs from pine seedlings was observed under a fluorescence microscope. After the observation of fungal infection, the PWNs were stored at −20 °C for the molecular qualification measurement of *E. vermicola*.

#### 4.4.4 Temporal and spatial colonization of *E. vermicola* in wilting pine trees

PWN-infected 10-year-old *P. koraiensis* trees showing wilt symptoms were used to reveal the colonization patterns of *E. vermicola* in infected trees. The pine trees grown in the greenhouse were infected with 20,000 PWNs via drill holes in the trunk. Subsequently, a blastospore suspension (3 × 10^8^ ml^-1^) of *E. vermicola* mutant *CNU120806gfp* was inoculated into pine tree xylem via injection at approximately 45 days after PWN infection when the symptoms of pine wilt disease appeared. Simultaneously, the PWN-infected wilting trees were treated with sterile distilled water for the control. The inoculation of *E. vermicola* followed the injection method described in the previous section. After that, the wilting pine trees were cut at 0, 7, 14, 21, 28, 42, 60, and 90 dai, and wood samples were stored at 4 °C. Three duplicates were conducted.

The wood samples within 50 cm of the inoculation holes were cut lengthways into several pieces at intervals of 5 cm from the inoculation hole in both the upward and downward directions for subsequent experiments. First, fungal colonization in wilting pine xylems was observed under a fluorescence microscope after free-hand sectioning. Meanwhile, the PWNs were extracted from the wood pieces using the Baermann funnel technique and measured under a white light and fluorescence microscope. The PWN densities were calculated from the number of isolated PWNs and the weight of wood samples. Subsequently, the wood samples and extracted PWNs were stored at −20 °C for total DNA extraction and TaqMan PCR quantification of *E. vermicola*.

### 4.5 Real-time PCR quantification of *E. vermicola*

#### 4.5.1 Total DNA extraction of wood and PWN samples

All of the sampled wood specimens were removed from the bark and surface-sterilized with 5% NaOCl for 1 min to avoid the influence of hyphae attached to the surface of the materials [38,39]. Then, the wood samples were ground into coarse powder and stored at −20 °C for subsequent DNA extraction. Three duplicates were used for each treatment. Total DNA was extracted from 5-day-old pure cultures of *E. vermicola* mutant *CNU120806gfp* grown on PDA, wood samples, and PWN samples using a modified CTAB extraction method. Total DNA was extracted in 5 ml of CTAB extraction buffer (100 mM Tris-HCl pH 8.4, 1.4 M NaCl, 25 mM EDTA, and 2% hexadecyl-trimethyl-ammonium bromide) at 70 °C for 3 hours (30 min for pure cultured fungal mycelium and PWNs). Then, the extracts were transferred to new 15-ml tubes and purified with 1 volume of phenol-chloroform-isoamyl alcohol (25:24:1) and with 1 volume of chloroform-isoamyl alcohol (24:1). Subsequently, the DNA was precipitated with 1 volume of absolute ethanol. DNA pellets were washed twice in 75% ethanol, dried in an air hood, and resuspended in 200 μl of 1 × TNE buffer (10 mM Tris-HCl pH 7.4, 200 mM NaCl, 1 mM EDTA). Then, ribonuclease A was added, and the samples were incubated for 30 min at 37 °C. The suspensions were purified with phenol-chloroform-isoamyl alcohol and chloroform-isoamyl alcohol and precipitated with 1 volume of absolute ethanol. Total DNA was resuspended in RNA-free water. The DNA samples extracted from fungal mycelium were visualized on 1% agarose gels stained with SYBR Green under ultraviolet light and quantified using spectrophotometry with a NanoDrop 2000 (Thermo Scientific, Pittsburgh, PA, USA).

#### 4.5.2 Quantification of *E. vermicola* in wood and PWN samples

The total DNA samples were extracted from wood and PWNs using the method described by Wang et al. (2020a) [18]. Then, the DNA extracts were used for quantification of *E. vermicola* using the TaqMan probe PCR technique to discover the fungal colonization pattern in the host pine tree and infection of PWN. In each reaction, 100 ng of total DNA extract was included as a DNA template to quantify the amount of target DNA. The *C*_*t*_ values obtained from TaqMan PCRs containing total DNA extracts of wood samples were interpolated using calibration curves to calculate the quantity of target DNA with respect to the total DNA.

### 4.6 Observation of fungal colonization in pinewood

To observe the fungal germination and colonization of *E. vermicola* in the host pine tree, wood inoculated with mutant *CNU120806gfp* was used for visualization by sectioning and fluorescence microscopy. The pine xylem of wood samples was free-hand sectioned with a slicing knife to obtain thin wood slices. Subsequently, the fragments of the sectioned wood slices were randomly picked and mounted on glass slides with a cover glass. Then, the wood slices were observed and recorded with white and fluorescence microscopy.

### 4.7 Statistical analyses

Data obtained from experiments were compared using ANOVA tests for differences between treatments using SPSS (version 20.0; SPSS, Chicago, IL, USA). The level of significance in all cases was 95%. Then, the statistical analyses were visualized by SigmaPlot (version 16.0; Inc., San Jose, California, USA).

## Acknowledgments

This work was supported by the National Key R & D Program of China (2018YFC1200400 and 2017YFD0600100) and the National Natural Science Foundation of China (31300543). In addition, this work was supported by the China Scholarship Council.

## Conflicts of interests

The authors declare no competing financial conflicts of interests.

